# Integrative multi-layer workflow for quantitative analysis of post-translational modifications

**DOI:** 10.1101/2025.01.20.633864

**Authors:** Cristina A. Devesa, Rafael Barrero-Rodríguez, Andrea Laguillo-Gómez, Victor M. Guerrero-Sánchez, David del Río-Aledo, Diego Mena, Inmaculada Jorge, Estefanía Núñez, Enrique Calvo, Juan Antonio López, Consuelo Marín-Vicente, Ana Martínez-Val, Emilio Camafeita, Borja Ibáñez, José Antonio Enríquez, José Luis Martín-Ventura, Jose Manuel Rodríguez, Jesús Vázquez

**Affiliations:** Centro Nacional de Investigaciones Cardiovasculares (CNIC), Madrid, Spain; CIBER de Enfermedades Cardiovasculares (CIBERCV), Madrid, Spain; CIBER de Fragilidad y Envejecimiento Saludable (CIBERFES), Madrid, Spain; Instituto de Investigacion Sanitaria Fundacion Jimenez-Diaz-Autonoma University of Madrid (IIS-FJD, UAM), Madrid, Spain

## Abstract

Novel algorithms based on ultratolerant database searching have paved the way for comprehensive analysis of all possible post-translational modifications (PTM) that can be detected by mass spectrometry-based proteomics, obviating their prior knowledge. These tools together with novel quantitative statistical models allow hypothesis-free approaches to study the role and impact of PTM on biological systems. However, interpretation of this information from a pathophysiological perspective is challenging due to the huge amounts of PTM data, the existence of chemical, structural, and statistical artifacts and the lack of dedicated tools for their analysis.

Here we propose a novel integrative workflow that automatically captures several layers of PTM-related information, including variations in trypsin efficiency, zonal changes, specific PTM changes and hypermodified regions, allowing advanced control of artefacts and coherent and comprehensive interpretation of PTM data. We show the performance of the new workflow by reanalyzing proteomics data from animal models of mitochondrial heteroplasmy and ischemia/reperfusion, revealing relevant PTM information not previously detectable, including consistent detection of novel oxidative modifications in Met and Cys residues from raw proteomics data. The workflow is available through the application PTM-compass.

## Introduction

Protein post-translational modifications (PTMs) significantly enhance proteome complexity and are essential for regulating protein localization, activity, folding, interactions, and stability. Consistent with their functional significance, PTMs have increasingly been recognized as playing a pivotal role in major diseases like cancer, as well as neurodegenerative and cardiovascular disorders ^1–3^. Owing to its specificity, sensitivity and high throughput capacity, mass spectrometry (MS) has become the gold-standard technique for the analysis of PTMs. However, MS has traditionally been applied using hypothesis-driven approaches, where the mass of the modification is known a priori. It was not only relatively recently that the so-called “open search” (OS) strategy ^4^ paved the way for true hypothesis-free analysis of PTMs. OS algorithms combine conventional search engines with peptide precursor mass tolerances of hundreds of daltons to fit any mass difference stemming from the modification. Later, some OS engines, such as Comet-PTM ^5^ or MSFragger ^6^, were improved to take into account the mass shift to match the fragments containing the modification.

Built upon our previous workflow aimed at the quantification of peptides and proteins ^7, 8^, we reported the first algorithm specifically designed for the quantitative analysis of posttranslationally modified peptides ^5^. In addition, the open-source iSanXot application ^9^, a recent implementation of the generic integration algorithm (GIA) ^8^, provided a user-friendly interface for adapting the above-described PTM quantitation workflow to fit any proteomics experiment. In spite of these developments, biological interpretation of the information obtained from high-throughput analysis of PTM remains challenging due to the huge amounts of PTM data produced, the existence of chemical, structural, and statistical artifacts and the lack of dedicated tools for their analysis.

In this work, we propose a shift in perspective on PTM quantitation. We leverage the versatility of iSanXoT to design an integrative quantification workflow that decomposes the quantitative information into separate layers. This approach is able to capture biologically-relevant PTM with improved specificity, discriminating them from artifactual changes, variations in trypsin digestion efficiency and zonal protein changes. The new integrative strategy revealed hypermodified regions and relevant PTM information not previously detectable. These analysis can be automatically performed using a Nextflow pipeline named nf-PTM-compass followed by iSanXoT analysis using a dedicated PTM quantification workflow.

## Results

### A peptide-centric integration pathway for the analysis of PTM changes dissects three independent modification events

In our previous workflow for the quantitative analysis of PTM, we proposed to integrate the information at the peptide level (p) to the protein level (q) (p2q integration), using the GIA algorithm (Fig. 1A). This integration models the variance and standardizes the quantitative peptide values around the corresponding protein average^5^. Here we propose to decompose the p2q integration introducing intermediate levels to account for confounding factors (Fig. 1B). To define these levels, we introduce a more specific nomenclature, where the term *peptidoform* (pdm) is used instead of *peptide* (p), since OS approaches often identify multiple modified forms of the same peptide. A peptidoform is specifically defined as a peptide p containing a modification of mass d in position m, following the notation of Fig. 1B. In the pdm2pgm integration, pdm elements are first grouped to pgm elements, where g is a user-defined group of modifications. This integration mainly serves to join peptidoforms that behave as non-modified (NM) peptides, and is justified for the importance that our workflow will later give to the behaviour of NM forms (see below).

**Figure 1.**
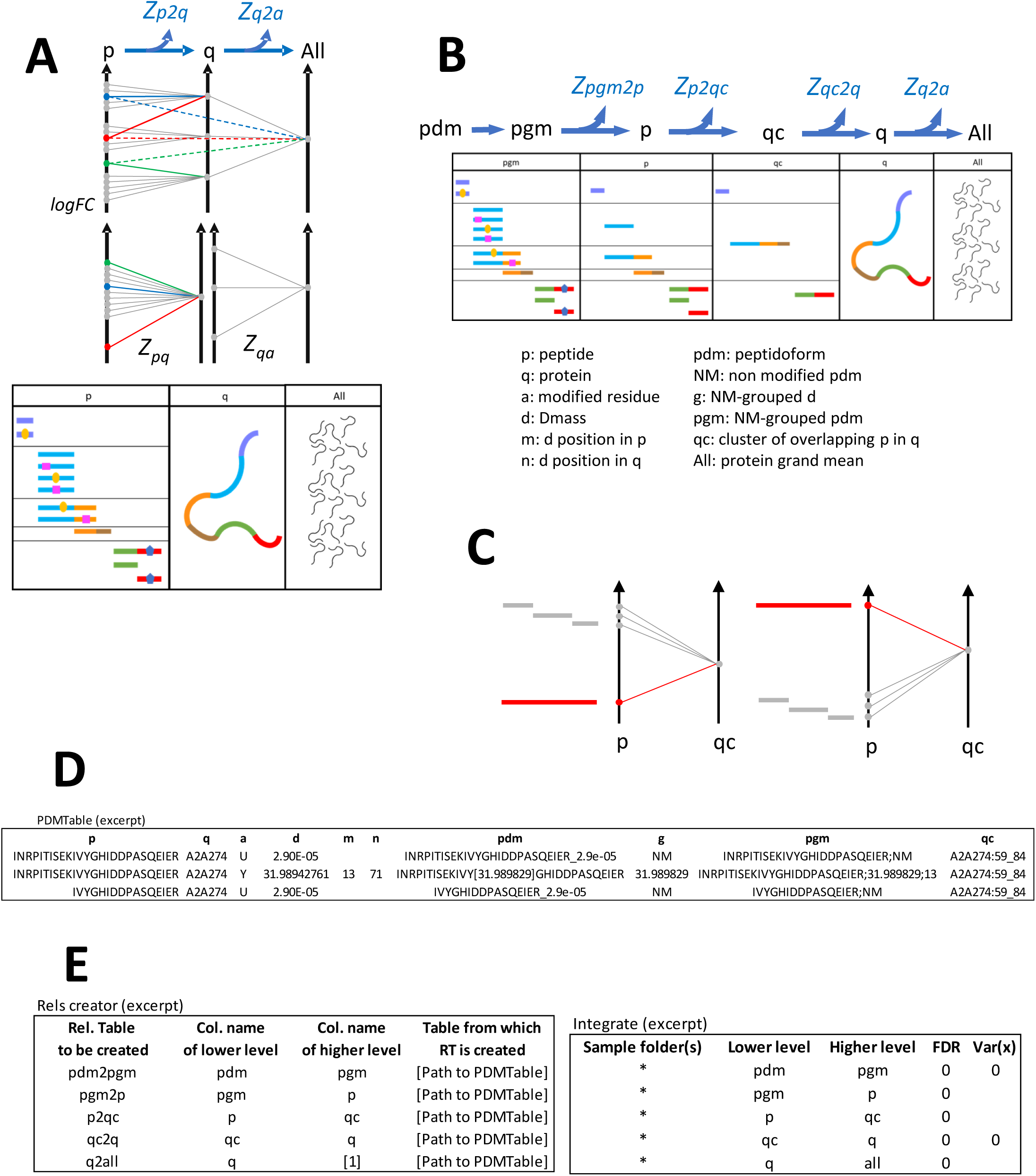
The peptide-centric integration pathway. A) Scheme of the original GIA workflow used for the quantitative analysis of PTMs. Note that integration to the protein level avoids false PTM changes (blue lines), detects true changes (red lines) and corrects as increases observed decreases (green lines). B) Scheme of the peptide-centric integration pathway, depicting the new integration levels. C) Scheme of the p2qc integration, which measure alterations in trypsin digestion efficiency. D) Excerpt from a PDM table showing elements of the different integration levels for two peptides with different peptidoforms that belong to the same cluster. E) Table depicting the information required by the two iSanXoT modules Rels creator and Integrate to carry out these integrations.

In the new integrative workflow, which we call the peptide-centric or p-pathway, pgm elements are firstly integrated to the p level (pgm2p). In this step, peptidoform changes are detected if they deviate from other peptidoforms originating from the same peptide (rather than from the same protein, as in the original workflow). Peptides (p elements) are then integrated into peptide clusters qc. A qc cluster is a region (or *zone)* of a protein q defined by starting and ending positions (defined in the element c) that contain a set of peptides partially overlapping due to partial digestion, so that clusters never overlap. The p2qc integration serves to measure the variability within the cluster and to detect peptide changes produced by variations in trypsin efficiency (Fig. 1C). Finally, qc clusters are integrated to the corresponding protein (qc2q). This last integration serves to measure variability across different regions of the protein and to detect zonal changes. To perform these integrations, we developed nf-PTM-compass, a Nextflow pipeline that automatically preprocesses the raw quantitative information in a format that can be directly analyzed using iSanXoT (see Methods). Nf-PTM-compass generates a so-called PDMTable that organizes pdm elements in rows and their properties in columns (Fig. 1D). This table is used by iSanXoT to automatically generate the relation tables needed and to perform the integrations as specified (Fig. 1E).

To test the novel workflow, we reanalyzed previously published proteomics data from studies on mitochondrial heteroplasmy in several mouse tissues ^5^ and ischemia/reperfusion (IR) injury in pig myocardium ^10^. The raw data was subjected to OS using MSFragger, preprocessed with nf-PTM-compass and analyzed using iSanXoT. We found that the variance of the pgm2p and p2qc integrations accounted together for the most part of overall variance in pdm2q, while that of pdm2pgm and qc2q was usually negligible (Fig. 2A). Consistently, when we tried to model the distribution of standardized values forcing the general variance to zero, we found that the GIA model could fit the distribution of Zpdm2pgm and of Zqc2q, but not the distribution of Zpgm2p or Zp2qc (Fig. 2B), clearly indicating the existence of an independent source of variability that affected the different peptidoforms derived from the same peptide or the different peptides pertaining to the same cluster. Furthermore, we were unable to find a significant correlation among the values of Zpgm2p, Zp2qc and Zqc2q (Fig. 2C). These findings support the notion that these integrations are reflecting truly independent events. In the absence of biological effects, clearly, the general variance of p2qc could be attributed to the technical variability introduced during trypsin digestion, while that of pgm2p to the manipulation of peptides before LC-MS/MS analysis.

**Figure 2.**
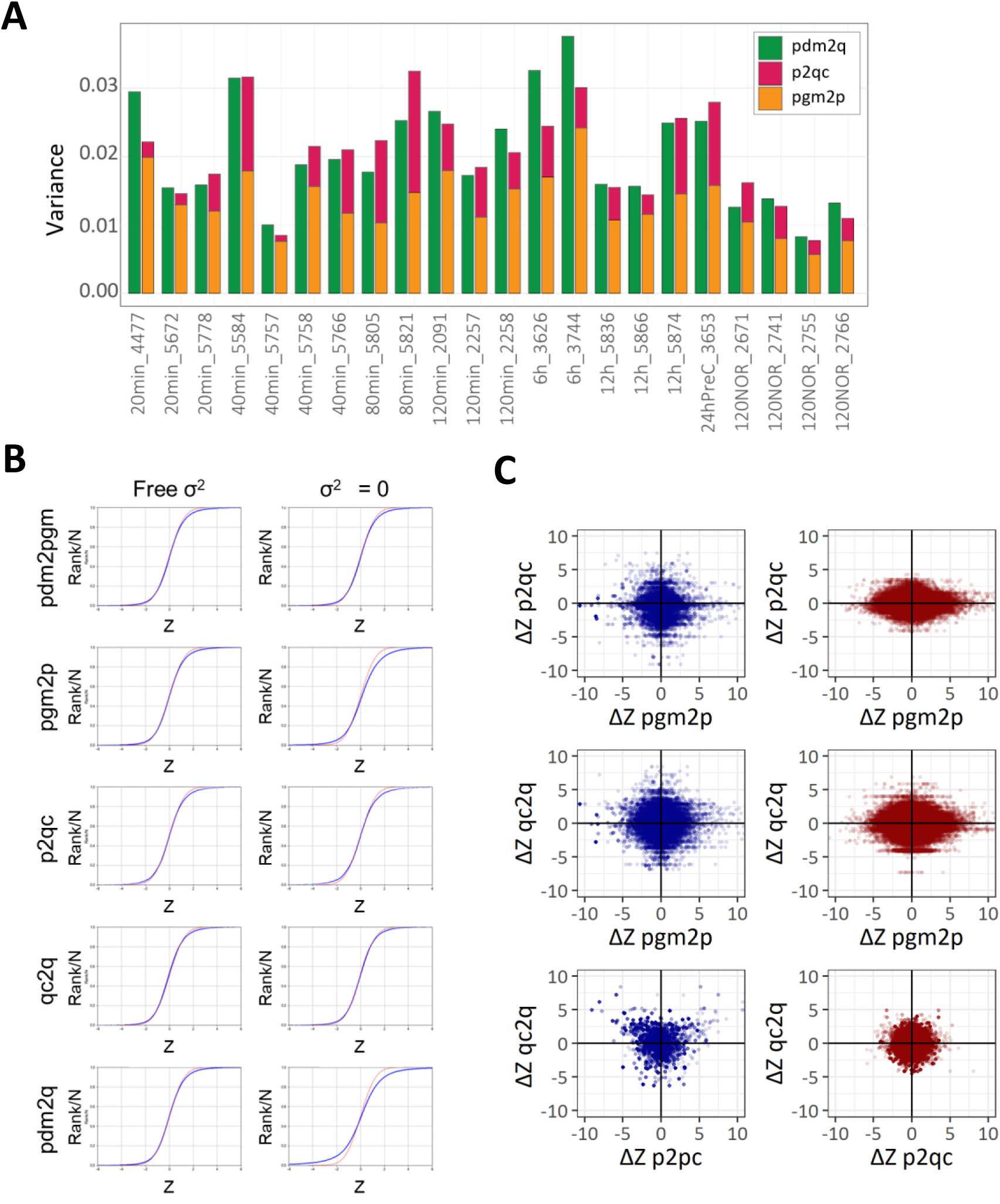
Error propagation across the different integration levels. A) Variance associated with the pdm2q (green), p2qc (purple), and pgm2p (orange) integrations for 22 cardiac tissue samples extracted from pigs subjected to varying reperfusion times. The variance derived from the intermediate qc2q integration was close to zero and therefore not represented. B) Distribution of Z, the standardized log2-ratio, for the different integrations considered in the peptide-centric pathway with unconstrained variance (left panel) or forcing variance to zero (right panel) in a sample of pig ischemic cardiac tissue. Note that in the latter case the pgm2p and p2qc integrations fail to fulfill the normal distribution, strongly suggesting that an additional source of variability affects the different peptidoforms derived from the same peptide or the different peptides pertaining to the same peptide cluster. C) Correlation between the difference in means of the Z values obtained from the pgm2p, p2qc, and qc2q integrations with mouse heteroplasmic cardiac tissue (left panel, blue dots) and pig ischemic cardiac tissue (right panel, red dots) supporting the independence of changes observed at each level from each other.

### The p-pathway identifies artefactual changes and delivers novel information

To evaluate the practical utility of the intermediate integrations included in the p-pathway, we compared the changes detected by pgm2p with those detected by pgm2q in heart tissue from heteroplasmic mice. Here we integrated pgm elements instead of pdm elements to eliminate the potential confounding effect of the pdm2pgm integration. As shown in Fig. 3A, top panel (red dots), a significant proportion of changes were detected by pgm2q but not by pgm2p. Inspection of these changes revealed that they were artefacts caused by variations in trypsin digestion efficiency, which affected all the peptidoforms derived from the same peptide and hence were false PTM changes (Fig. 3B, Zone A). Furthermore, we found a proportion of changes detected by pgm2p that were not detected by pgm2q (Fig. 3A, blue dots). These changes corresponded to elements that follow a pattern of change different from the rest of peptidoforms of the same peptide and were thus true PTM changes that were overlooked by pgm2q (Fig. 3B, Zone B). Similar results were obtained when analyzing results from the IR pig model (Fig. 3A, bottom panel).

**Figure 3.**
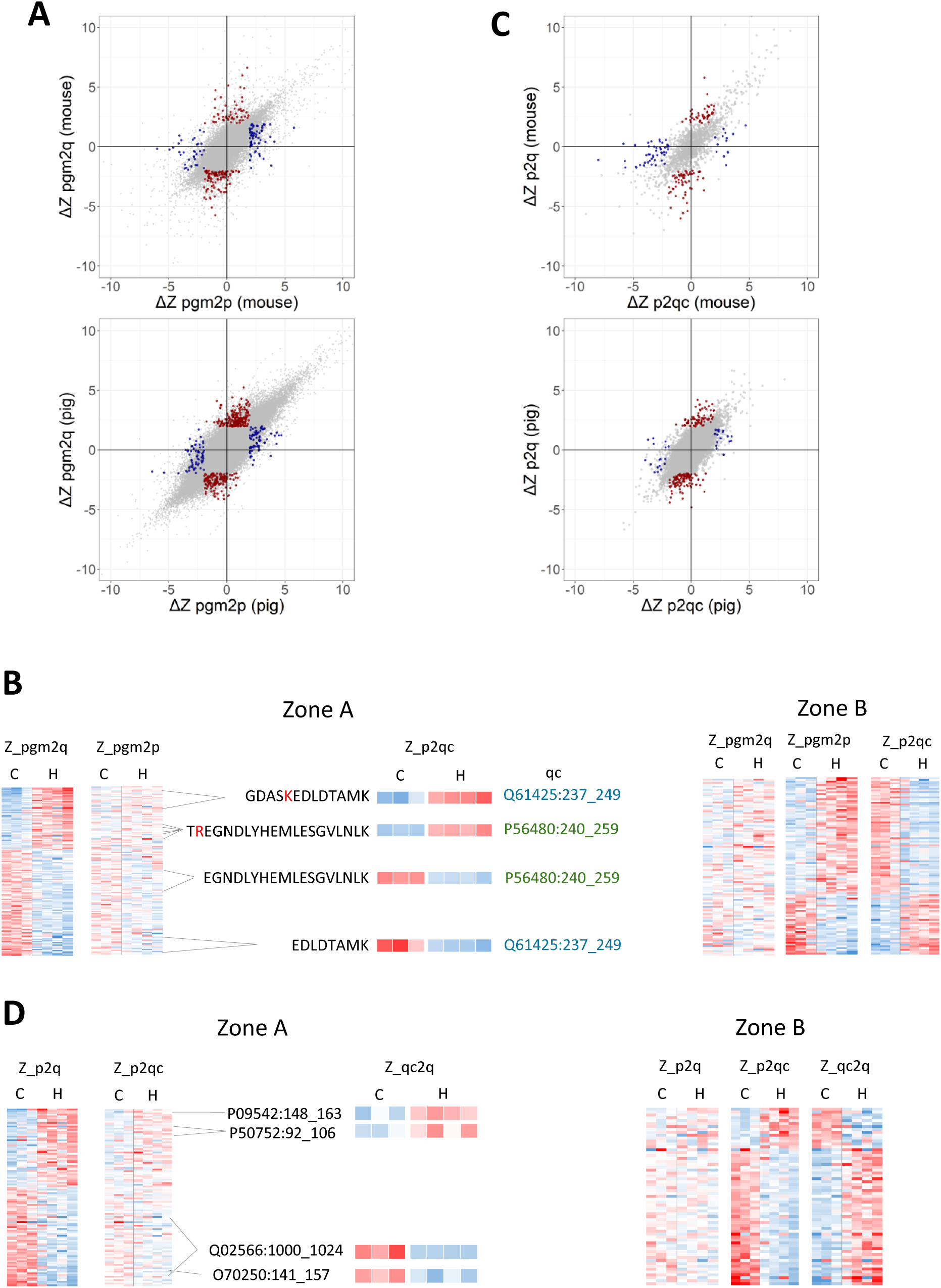
The peptide-centric pathway resolves artifacts derived from trypsin digestion efficiency and zonal changes. A) Correlation between the difference in means of Zpgm2p and Zpgm2q from mouse heteroplasmic cardiac tissue (top) and pig ischemic cardiac tissue (bottom) showing alterations revealed exclusively by pgm2q (red dots) or pgm2p (blue dots). B) Data taken from heteroplasmic mice heart showing how pgm2q changes undetected by pgm2p are artifacts originated by an alteration in trypsin digestion efficiency (left panel), and how changes detected only by pgm2p correspond to true PTMs (right panel). C) and D) Same plots as in A) and B) showing how p2q may produce peptide artifacts that are not detected by p2qc, and how p2qc detect true peptide changes that are not detected by p2q.

We then compared the results obtained by applying p2qc, the second integration of the p-pathway, with the direct integration p2q. Again, we found changes detected by p2q that were not detected by p2qc (Fig. 3C, red dots). These were artifactual changes that affected all the peptides from the cluster and were actually a consequence of a zonal change (Fig. 3D, Zone A). Finally, we also found changes detected by p2qc that affected differentially the peptides pertaining to the same cluster due to altered trypsin digestion efficiency and were not detected by p2q (Fig. 3F, blue dots, and Fig. 3D, Zone B).

### Trypsin digestion efficiency and zonal artifacts are reproducible within the same biological context

The ability to detect artifacts related to trypsin digestion efficiency and/or zonal changes raises the question of whether these artifacts are related to the biological setting in which they are observed. To answer this question, we compared the p2qc and qc2q changes observed in heart with those found in liver from heteroplasmic mice, two kind of samples that were prepared simultaneously during the same experiment but came from tissues that were known to behave in a different manner in response to heteroplasmy^11^. We found no evidence of correlation between them (Fig. 4A, left). Likewise, no correlation was observed for these variables when comparing the changes detected during the first oxidation wave (20 min) with those detected during the second wave (24 h) in the IR study in pig, two waves shown to be produced by different biochemical events^12^ (Fig. 4A, right). In clear contrast, when we compared p2qc changes obtained with hearts from different groups of pigs within the second oxidation wave, we found a clear correlation between them (Fig. 4B). In fact, the pattern of changes in trypsin digestion efficiency were, in general, highly reproducible within the same wave and protein (Fig. 4C).

**Figure 4.**
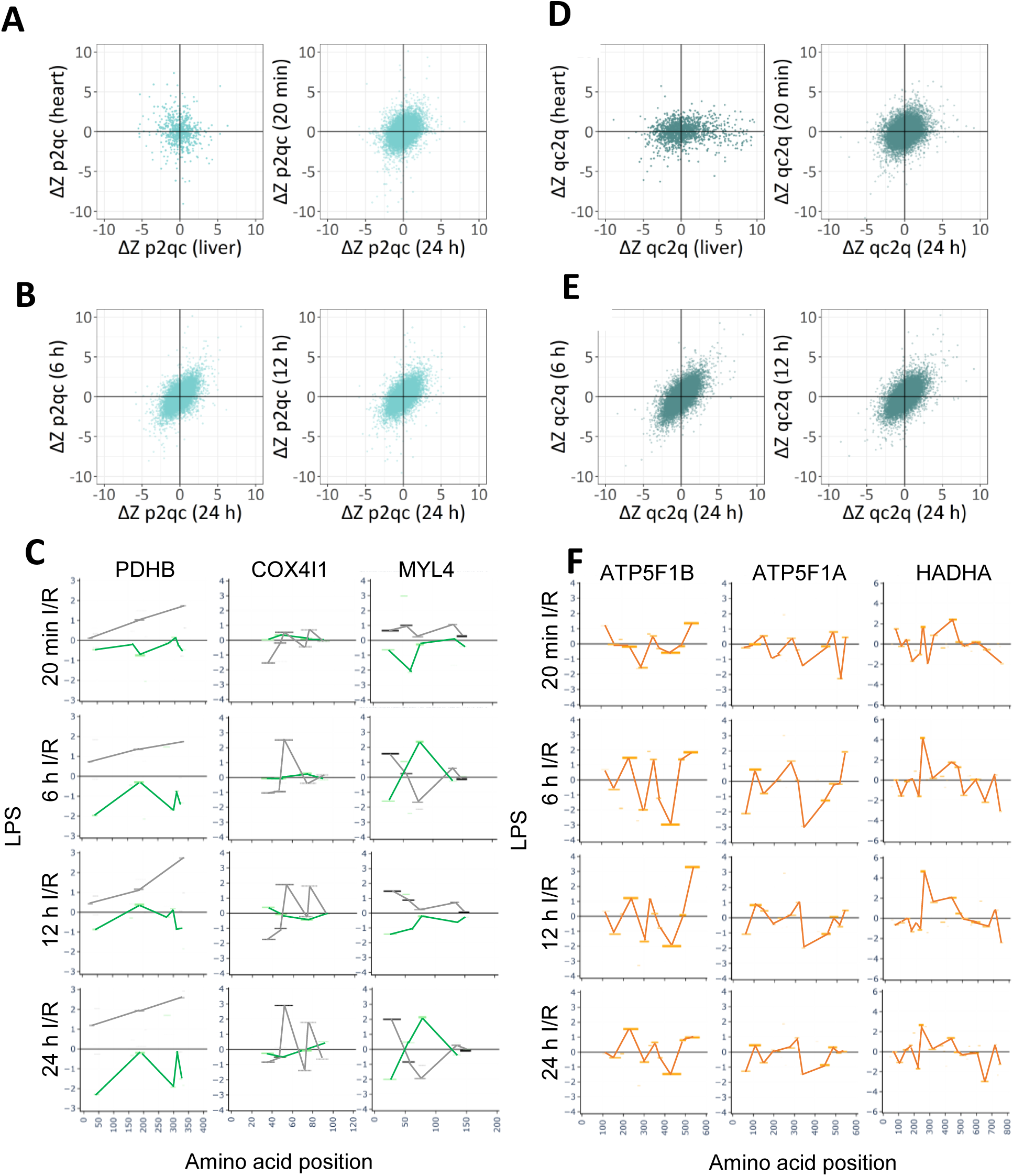
Reproducibility of trypsin accessibility and zonal changes. A) Correlation between the difference in means of Zp2qc from mouse heteroplasmic liver and heart tissues (left) and from pig heart tissue collected at two separate oxidation waves after ischemia/reperfusion (right). B) Correlation between the difference of means of Zp2qc from pig heart tissue collected within the same IR oxidation wave (left: 24 h vs 6 h reperfusion time; right: 24 h vs 12 h reperfusion). C) Representative examples of p2qc maps showing the reproducibility of trypsin efficiency changes in three selected proteins from pig heart tissue collected at different times after iR; note that 20 min corresponds to the first wave and 6, 12 and 24 h to the second wave. The X-axis is the amino acid position in the protein; LPS in the Y-axis represents the co-logarithm of t-test p-values comparing means between conditions multiplied by the sign of the change (up: increases, down: decreases). The green and black lines join totally and partially digested peptides, respectively. D) and E) Same plots as in A) and B) correlating Zqc2q values between different samples. F) Representative examples of qc2c maps showing the reproducibility of zonal changes in three selected proteins, as in C). The yellow lines join the zonal changes.

Similar results were found when qc2q values were compared between different samples. Zonal changes were quite dissimilar between samples from different heteroplasmic mice tissues or from the heart of pigs from different oxidation waves (Fig. 4D). In clear contrast, zonal changes showed a highly reproducible pattern within the same oxidation wave and protein. (Fig. 4E and F). These results strongly support the idea that trypsin efficiency and zonal changes are artifacts closely related to the biological nature of the samples analyzed and highlight the importance of identifying them to prevent their interference with the detection of genuine PTM changes.

### The p-pathway allows detection of specific PTM changes and hypermodified regions

While inspecting the results generated by the p-pathway workflow, we found an additional source of artifacts that needed further statistical refinement. When pgm2p integrates peptidoform quantifications into the peptide value, PTM changes are expected to be detected as a significant deviation from the averaged peptide value (Fig. 5A, left). However, in a large number of cases we found that the NM peptidoform, together with other modified peptidoforms (which we call non-specific PTMs) and related artefacts, showed changes in the opposite direction that became statistically significant (Fig. 5A right). To avoid this effect, we used the NM peptidoform as a reference to determine which changes were specific. This was accomplished by subtracting the value of the corresponding NM peptidoform from the Zpgm2p value of each peptidoform in each sample, thereby obtaining an NM-corrected value (Zpgm2p_NM, Fig. 5B, upper and medium panels).

**Figure 5.**
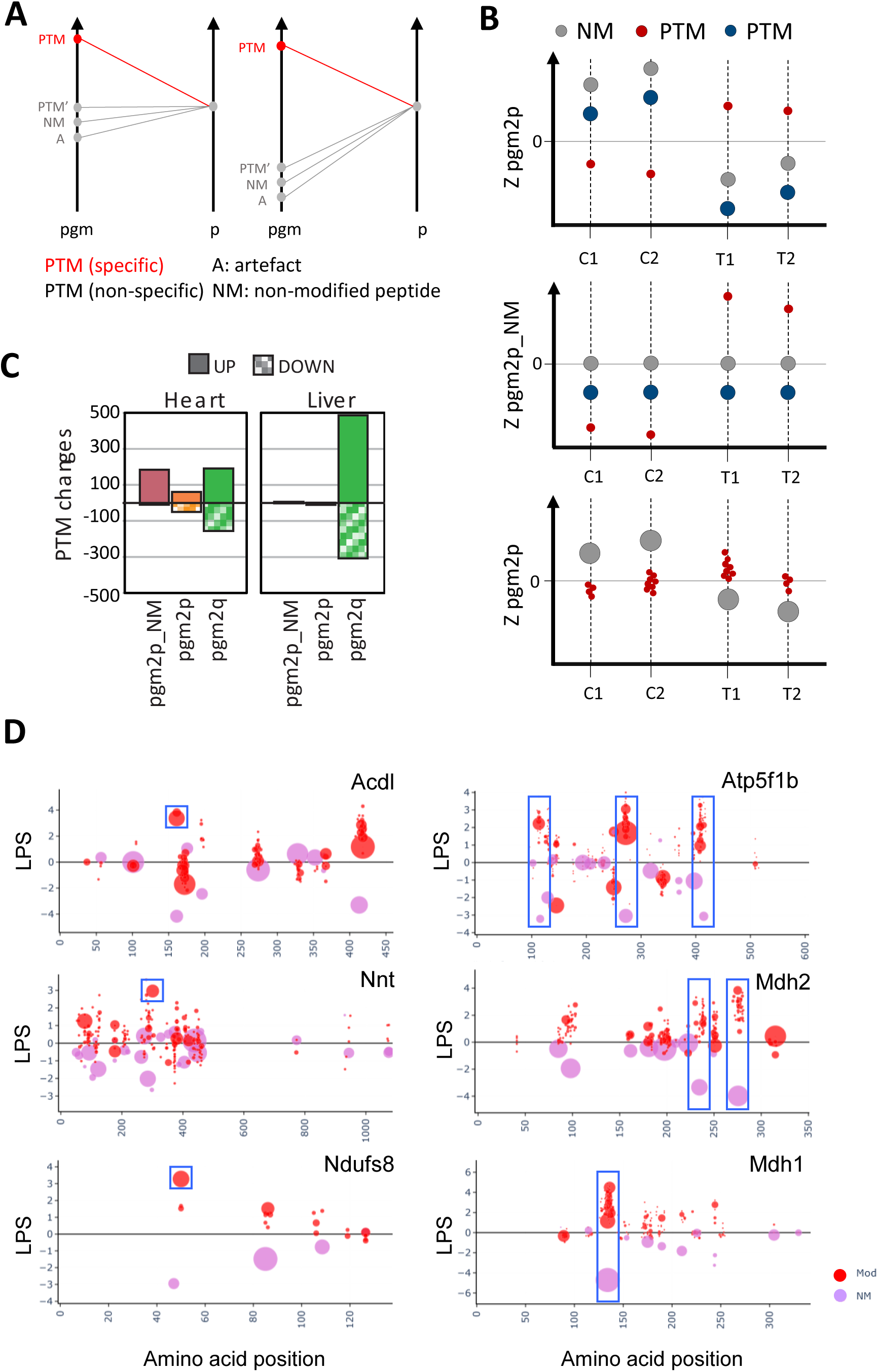
Detection of specific PTM changes and of hypermodified regions. A) Scheme of the pgm2p integration used to assess the specificity of PTM changes. Specific PTM changes are detected as significant deviations from the peptide average (left); however, the NM peptidoform, together with other modified peptidoforms and related artefacts, can also deviate statistically significantly from average in the opposite direction (right). B) Scheme of the strategy adopted to detect specific PTM changes using the NM peptidoform as a reference (upper and medium panels). C and T represent control and treated samples, respectively. The pgm2p integration can also reveal hypermodified regions harboring heterogeneous PTMs by detecting changes in NM peptidoforms (lower panel). C) Number of significant changes observed in the indicated integrations in mouse heteroplasmic heart and liver tissue. D) Examples of specific PTM changes (left panel, squared dots) and hypermodified regions (right panel, squared areas) detected in selected proteins from heart tissue from heteroplasmic mice. The X and Y axis are as in Fig. 4D and F. The red dots show LPS values from modified forms, calculated by Zpgm2p_NM, and the magenta dots the LPS values from NM peptidoforms, calculated by Zpgm2p. The size of the dots are proportional to the abundance of the peptidoform in spectral counts.

To analyze the practical impact of using NM-corrected values, we compared the results obtained with or without applying the specificity criterion in the heteroplasmy model. By directly integrating the peptidoforms to the protein value (pgm2q), we observed significant increases and decreases in both heart and liver from heteroplasmic animals (Fig. 5C, green bars). These changes were not observed in liver when integrating to peptide instead (pgm2p), but remained symmetrically distributed in heart, showing increases and decreases (Fig. 5C, brown bars). In clear contrast, by applying the specificity criterion using Zpgm2p_NM values, the majority of significant changes in heart were peptidoform increases (Fig.5C, magenta bars), in agreement with the concept that heteroplasmy induces protein modifications by oxidative damage specifically in the heart, a tissue unable to resolve heteroplasmy, but not in the liver^11^.

We then investigated whether this approach was able to detect specific Met oxidation changes. This modification is often considered a non-specific artifact produced by spontaneous oxidation; however, numerous publications have demonstrated that Met oxidation may be produced in specific biological contexts, often playing regulatory roles on protein functionality. Taking the data from pgm2q, we found that 51% of changes were observed in peptidoforms whose affected Met site was only observed in the mono-oxidized form, as happens when Met is oxidized during sample preparation. In contrast, in the vast majority of changed peptidoforms (75%) detected by applying the specificity criterion (i.e. using pgm2p_NM values), the oxidation affected a Met residue that was observed to harbor other modifications. This suggests that Met oxidation changes detected by pgm2p_NM were not artifactual, but the first step in a process of multiple oxidative events produced in specific Met residues in a context of oxidative stress.

To further explore this effect, we constructed *protein PTM-maps* by plotting the change in the abundance of all observed peptidoforms for a given protein versus their amino acid position in the protein sequence. We noticed several cases where the specific PTM changes were unique or the predominant modification in the peptide (Fig. 5D, left, red dots within blue squares). However, in approximately half of the cases, we also observed groups of altered peptidoforms that concentrated around certain amino acid positions, which we called “hypermodified regions” (Fig. 5D, right, blue rectangles). These regions contained numerous modified peptidoforms whose abundance changes occurred in the same direction, while the corresponding NM peptidoform exhibited the opposite change (Fig. 5D, purple balls). On the basis of these observations, we decided to perform a separate analysis by searching for statistically significant changes in Zpgm2p values from NM peptidoforms only, to systematically detect the presence of hypermodified regions (Fig. 5B, lower panel). We detected a total of 49 hypermodified regions in heart (Fig. 6A, left). Of these, the large majority (45) corresponded to decreased NM peptidoforms, thus reflecting the homogeneous tendency to increase for the 1,751 different modified peptidoforms accumulated in these regions (Fig. 6A, right).

**Figure 6.**
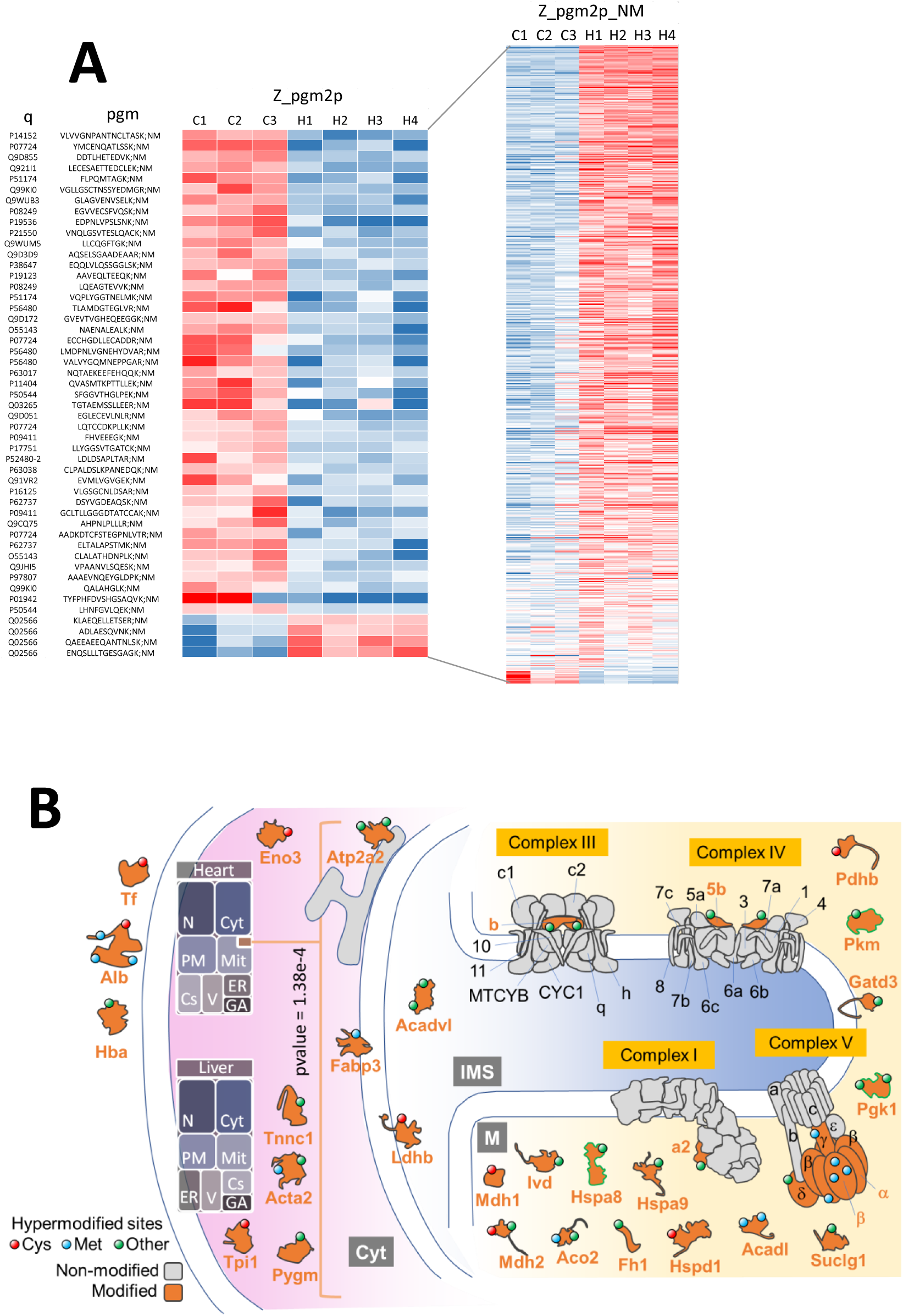
Systematic detection of hypermodified regions. A) Zpgm2p values from NM pgm elements with statistically significant change in heteroplasmic heart mice mice (left panel) and Zpgm2p_NM values from all the modified pgm that belong to the same peptides (right). B) Map of proteins harboring hypermodified regions. The circles indicate the main modified residue (blue - Met, red - Cys, and green - other residues). The treemap boxes indicate the number of proteins from N – Nucleus, Cyt – Cytosol, PM – Plasma Membrane, Mit – Mitochondria, Cs – Cytoskeleton, V – Vesicles, ER – Endoplasmic Reticulum, and GA – Golgi Apparatus. Green bordered proteins have been described as mitochondrial under stress conditions. M, mitochondrial matrix; IMS, inter membrane space.

This method for systematic detection of hypermodified regions allowed us to construct a map of hypermodifications in heteroplasmic heart (Fig. 6B). We found that the proteins containing hypermodified regions were significantly enriched in mitochondria, where they concentrated in the mitochondrial matrix. These included proteins from the matrix-oriented moiety of the electron transport chain complexes, being Complex V the most affected, and also from the TCA cycle and fatty acid and carbohydrate metabolism.

### Hypermodified regions display consistent patterns of oxidative modifications on Met and Cys sites

To analyze the nature of the hypermodified regions detected in the heart from heteroplasmic mice, we selected the most abundant peptidoforms thereof and examined the spectral count frequency with which each amino acid was found modified. We found that the majority of hypermodified regions were located around Met and Cys residues (Fig. 7A).

**Figure 7.**
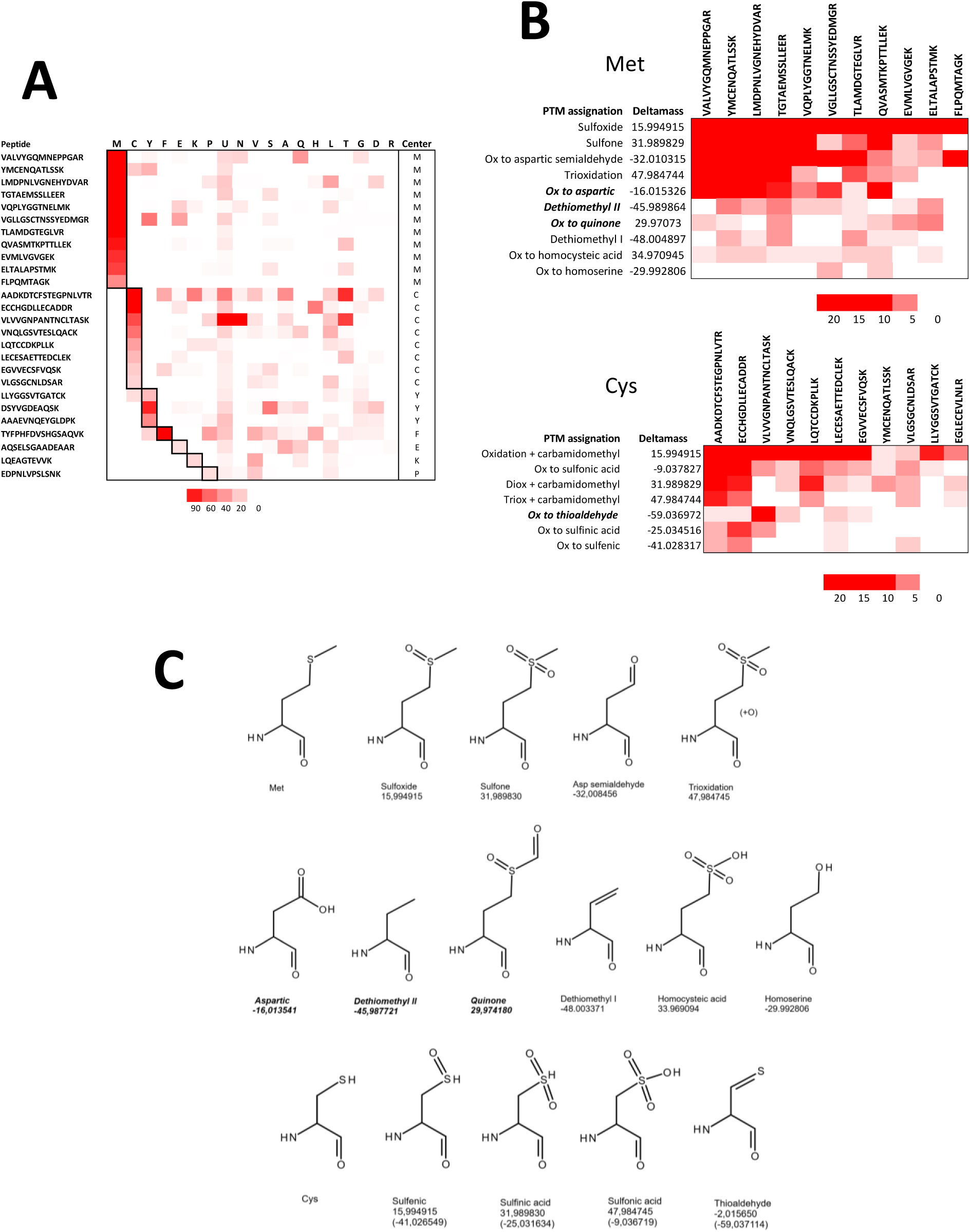
Analysis of hypermodified regions in heteroplasmic mice and detection of novel modifications in Met and Cys. A) Heatmap showing spectral count abundance of the amino acids bearing modifications in the hypermodified regions. The column *Center* indicates the most probable residue hipermodified. The table contains hypermodified regions with more than a total of 45 spectral counts. B) Heatmap showing spectral count abundance of ΔMass values across the hypermodified regions centered in Met (top panel) and Cys (bottom panel) residues. These tables are filtered for ΔMass values that are found in at least two different peptides with at least two spectral counts. C) Tentative chemical structures of some of the oxidative Met and Cys modifications specifically increased in heart from heteroplasmic mice. The names of the structures not described in the Unimod database are highlighted in bold italics. The numbers indicate the theoretical ΔMass values.

Interestingly, hypermodified Met residues showed a tendency to harbor the same set of oxidative modifications (Fig. 7B, top panel). Besides sulfoxide (mono-oxidation), Met was found in sulfone form (dioxidation) and also in eight extra states where the observed mass shift agreed well with tentative additional oxidative forms (Fig. 7B, top panel, and 7C). These included oxidation of Met to aspartic semialdehyde, homocysteic acid and homoserine, as well as a form derived from the loss of the oxidized thiomethyl moiety, as described in the Unimod database. A trioxidized Met form was also assigned, although it probably corresponds to the oxidation of adjacent amino acids. Strikingly, the mass shifts were also in good agreement with tentative Met oxidation to aspartic acid and quinone forms, and with the loss of the thiomethyl side chain from oxidized Met by a different route, none of which are described in Unimod (Fig. 7B, top panel, and 7C; italics, bold characters).

Inspection of the hypermodified forms centered in Cys revealed the widespread presence of the sulfinic and sulfonic acid forms and, in lower proportion, of the sulfenic acid (Fig. 7B, bottom panel, and 7C). Interestingly, we also found a form whose mass shift fitted well with a tentative oxidation of Cys to thioaldehyde (Fig 7B, bottom panel, and 7C). Taken together, these findings provide consistency and support the validity of the integrative workflow for improved detection of biologically-relevant modifications.

## Discussion

In previous works we proposed the WSPP model, a statistical framework for the quantitative analysis of proteins from MS-based bottom-up proteomics where the quantitative information at the peptide level was integrated to the protein level using an approach that modeled peptide variance ^13, 14^. This concept was later applied to the study of reversible Cys modifications by redox proteomics, where statistically significant changes in these PTMs could be detected as deviations from the modeled distribution of the rest of peptides ^15^. The model was generalized in the form of the Generic Integration Algorithm (GIA) ^8^, which could be applied using SanXoT, a dedicated software package ^16^. These tools have been employed to study redox PTMs in several biological contexts ^17–21^. We later reported that the GIA model could be used as a statistical framework for the quantitative, high-throughput analysis of all the PTMs detected following open-search approaches ^5^. The successful application of the GIA for the non-targeted analysis of PTMs in numerous biological contexts ^5, 10, 11, 21–27^situations showed the robustness and validity of this algorithm as a statistical framework for the quantitative analysis of PTMs.

The GIA employs relative abundance ratios as quantitative values and models the individual peptidoform variance (to which we refer as peptidoform-to-protein variance) as a deviation from the averaged protein value. This approach serves to standardize peptidoform values (i.e. to express them in units of standard deviation), so that they can be analyzed using a N(0,1) normal distribution as null hypothesis. The GIA thus analyzes peptidoform changes independently, so that they are not affected by protein changes. The development of iSanXoT, a standalone application that incorporates a user-friendly interface to freely define integration levels ^9^, has brought the GIA approach within the reach of the proteomics community.

In this work we leverage the flexibility provided by iSanXoT to show that the peptidoform-to-protein variances can be decomposed in at least three separate components generated by quantitative variations in: i) the different peptidoforms from the same peptide, ii) the different peptides from the same peptide cluster, and iii) the different peptide clusters from the same protein. We demonstrate that each of these components can be modeled using the GIA, that they vary independently from each other, and that they need to be considered to prevent misassignment of false PTM changes. These factors were identified from the observations of PTM changes in previous works that by their nature were expected to be artefacts. The first factor (i.e. peptidoforms within the same peptide) may be interpreted as the conventional variance at the peptidoform level, which is captured with improved specificity by integrating to the peptide rather than to the protein level. The second factor is generated by variations in trypsin digestion efficiency, and therefore may be considered as an undesired artifact, since it is well known that digestion efficiency is never complete in bottom-up proteomics. The third factor, herein referred to as zonal changes, may be tentatively explained by other events such as variations in the accesibility of trypsin to different protein regions, the existence of strong protein interactions or alterations to PTMs that are difficult to be detected by MS, such as glycosylation or binding to large lipid moieties. Importantly, regardless of their artifactual nature, our results indicate that the latter two factors are reproducible even in situations of large biological variability such as those analyzed in this work, where each biological replicate comes from tissues from different animals. These observations suggest that these factors are also reflecting biological variations, a possibility that will be cleared up in future studies. In any case, our approach allows a full control over these artifacts, considerably reducing the information that has to be interpreted and providing a much more precise representation of the pattern of PTM alterations.

The integration of peptidoform to peptide implies a normalization at the peptide level, whose quantitative value is calculated as an averaged mean of all its peptidoforms. This approach allows an accurate quantification of relative peptide abundances, as it makes use of all the quantitative information experimentally available, and allows appropriate integration of the information to higher levels. However, it also produces artifactual peptidoform changes, because PTM increases or decreases produce opposite changes in the rest of peptidoforms of the same peptide. This effect is easily corrected by using the non-modified peptidoforms as a reference for the detection of what we call *specific* PTM changes. We show here that this correction is effective to remove artifactual changes and further reduce the number of changes that need to be biologically interpreted. ^5, 11^

An interesting observation derived from the new workflow is that a significant proportion of specific PTM changes are not isolated, but take place in residues that are affected by other modifications. These PTM changes, concentrated in what we call *hypermodified regions*, can be readily observed from the inspection of the *PTM maps* of the affected proteins, which display how PTM alterations distribute along protein sequences. We propose analyzing the behaviour of the non-modified peptidoforms as a systematic method to detect hypermodified zones. This approach considerably reduces the number of elements to be analyzed, relaxing the constraints imposed by multiple hypothesis testing. Application of this strategy to the mice model reveals specific protein regions that concentrate PTM produced by heteroplasmy in heart. These regions accumulate mostly in proteins from the mitochondrial matrix and from the moiety of complexes of the electron transport chain that are oriented towards the mitochondrial matrix. These include several subunits of ATP synthase (Atp5f1a, Atp5f1b Atp5f1c and Atp5f1d) and cytochrome c oxidase (Cox5b) complexes, of Complex III (Uqcrb) and also Mdh2, integral to both the TCA cycle and the respiratory chain. Hypermodified regions are also found in proteins such as Acadl, Acadvl and Fabp3, which are implicated in lipid metabolism and transport, Atp2a2 and Tnnc1, which are implicated in calcium regulation, and Hspa8 and Hspa9, which belong to the HSP70 family and are implicated in stress and immune responses, among others. These data extend results found in previous analysis ^5, 11^, and suggest a potential effect of hypermodifications on mitochondrial bioenergetics, calcium handling, oxidative stress and inflammation. Notably, these regions were not detected in other tissues such as liver, that are able to resolve heteroplasmy.

The hypermodified regions detected in the heteroplasmic heart using this method were predominantly located around Met and Cys sites. Strikingly, analysis of these regions revealed the presence of oxidative Met modifications beyond the well-known Met sulfoxide that are consistently found in practically all of them. Thus, Met sulfone and other oxidized forms of Met described in the literature such as Asp semialdehyde ^28^ and homocysteic acid ^29^ were detected. We also found a trioxidized form, which may potentially arise from oxidation of adjacent amino acids, as well as a form typically produced by neutral loss of the oxidized thiomethyl moiety from Met sulfoxide upon collision-induced dissociation (dethiomethyl I) ^30^. Interestingly, we also found other Met modifications, not described in the Unimod database, that could be tentatively assigned to the oxidation of Met to aspartic acid, to quinone and to homoserine. While the oxidative conversion of Met to Asp has been described to take place in proteins ^31^ and Met has also been described to be converted to homoserine in the presence of strong oxidants like cyanogen bromide ^32^, to the best of our knowledge no previous evidence of the generation of carbonyl groups or the formation of homoserine from the oxidation of Met in biological conditions has been reported before. Finally, our results also suggest that Met oxidation may also produce a novel dethiomethylated form (dethiomethyl II) that has not been described previously. The hypermodified regions centered in Cys contained a mixture of the well-known sulfenic, sulfonic and sulfinic forms. The former two were also found in their corresponding carbamidomethylated adducts, expected from the iodoacetamide treatment ^33^. We also found an oxidized form not described in Unimod that could be tentatively assigned to Cys thioaldehyde and that has been observed upon peptide fragmentation of disulfide-linked peptides ^33^, but not directly in proteins. While the exact structure of these newly observed species deems further research, it is important to highlight that their systematic identification was made possible by the new workflow proposed here from raw LC-MS/MS data and without the need of enrichment procedures, whereas they were completely overlooked by previous quantitation methods ^5, 11^. Furthermore, these detailed patterns of consistent hypermodifications were only observed in heart, but not in liver or muscle from heteroplasmic mice, supporting their biological specificity.

The analysis of Met oxidation has garnered increasing interest within the proteomics community, as this modification is known to cause proteins to misfold and has been linked to a number of diseases, including neurodegenerative disorders and pathological aging ^34–37^. Met oxidation may modulate cell function through an interplay between specialized oxidases and sulfoxide methionine reductases that catalyze the oxidation and reduction of specific protein targets, respectively^34, 38^. However, measuring the levels of *in vivo* Met oxidation by MS-based proteomics is challenging due to the spontaneous oxidation of Met during sample processing and ionization and MS analysis of peptides^35, 39^. While some protocols have been proposed for the specific analysis of oxidized Met residues^35, 37, 40–42^, they are based on oxidative treatments or strong alkylating conditions that may modify other residues, thus precluding their general use in the context of unbiased PTM research.

Finally, one of the advantages of the novel integrative workflow is that it can be applied to any non-targeted LC-MS/MS data previously obtained. The systematic reanalysis of proteomics datasets using this new integrative concept may reveal novel biologically-coherent trends in the behaviour of PTMs and allow the discovery of novel modifications.

## Methods

The workflow has been implemented in a pre-processing Nextflow ^43^ pipeline, named nf-PTM-compass, which is used together with a novel iSanXoT workflow specifically designed for PTM quantification. An overview of the entire workflow is presented in Fig. 8.

**Figure 8.**
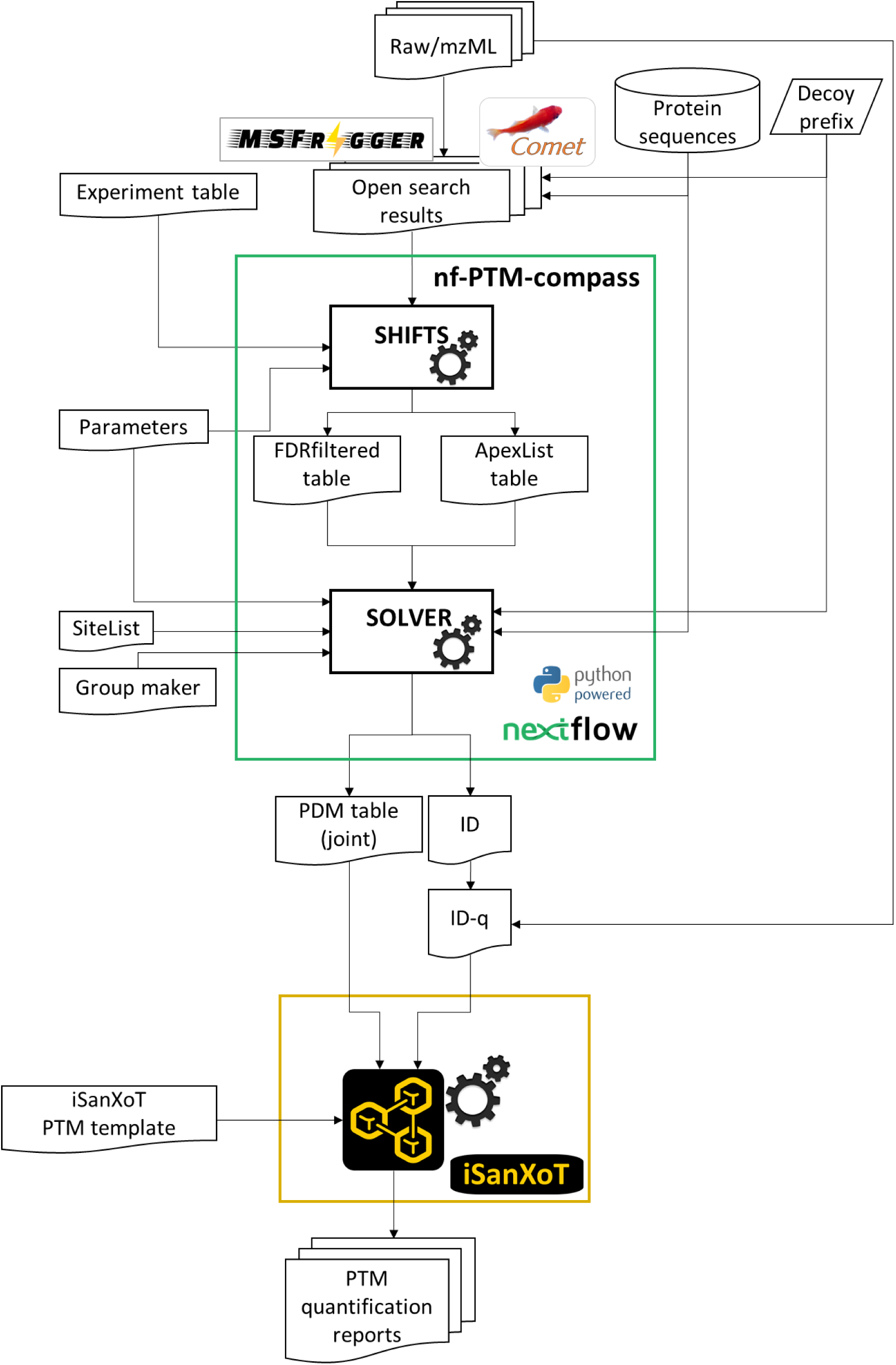
Overview of PTM-compass. Open search results from both Comet-PTM and MSFragger search engines are pre-processed using nf-PTM-compass, a pipeline implemented in Nextflow. Nf-PTM-compass generates a PDMtable containing the elements required by iSanXoT to perform the required integrations. Finally, the quantitative variables obtained for the different integrations carried out are reported in tabular form by iSanXoT.

The PTM-compass workflow accepts as input open search results from Comet-PTM ^5^ and MSFragger ^6^ search engines. The SHIFTS modules included in the nf-PTM-compass pipeline identify peaks in the ΔMass distribution, assign PSMs to peaks, and calculate the FDR for peptidoform identification, whereas the SOLVER modules are used to minimize peptidoform dilution and to generate the PDMTable. This table define the hierarchy of the elements that will be later used by iSanXoT (Fig. 1D). These pdm elements are then automatically processed by iSanXoT, which generates the relational tables required and performs the workflow integrations as specified (Fig. 1E).

nf-PTM-compass was developed in Nextflow ^43^, a framework specifically designed to create reproducible, scalable, and portable analysis workflows ^44^. In this implementation, the Nextflow pipeline integrates the Python scripts for SHIFTS and SOLVER, which are hosted in the PTM-compass repository. The SHIFTS and SOLVER scripts, along with their dependencies, are packaged in a Singularity container ^45^, ensuring streamlined deployment and reproducibility. The Nextflow pipelines can be executed on a variety of computing platforms, including Kubernetes clouds, local computers, and cluster environments such as SGE and others.

The nf-PTM-compass Nextflow pipeline is open-source and available in the GitHub repository: https://github.com/CNIC-Proteomics/nf-PTM-compass. Additionally, the Python scripts for SHIFTS and SOLVER can be found in the PTM-compass repository at https://github.com/CNIC-Proteomics/PTM-compass. To test the nf-PTM-compass pipeline, we provide result files obtained from the open search conducted using MSFragger ^6^ on the data reported in the heteroplasmy mice model study ^5^. For detailed instructions to execute this use case, please refer to the GitHub repository’s documentation.

The open-source iSanXoT ^9^ application provides a user-friendly interface, enabling the versatile and highly automatable creation of integrative quantification workflows and report tables. To adapt the PTM quantification workflow described above, we have provided an iSanXoT workflow template that generates the required integrations. This template includes the task tables necessary for each iSanXoT module involved in the PTM-compass workflow.

The iSanXoT workflow template, along with the required input files for executing this PTM-compass workflow, can be downloaded from: https://raw.githubusercontent.com/CNIC-Proteomics/iSanXoT/master/docs/templates/PTM-compass.zip. For detailed instructions on importing the workflow template, refer to the iSanXoT documentation at https://cnic-proteomics.github.io/iSanXoT/.

## Acknowledgements

This study was supported by competitive grants PID2021-122348NB-I00 funded by MICIU/AEI/10.13039/501100011033 and by “ERDF A way of making Europe”, PLEC2022-009298, PLEC2022-009235 and EQC2021-007053-P funded by MICIU/AEI/10.13039/501100011033 and by “European Union NextGenerationEU/PRTR”, and S2022/BMD-7333-CM (INMUNOVAR-CM) funded by Comunidad de Madrid. The project leading to these results has received funding from “la Caixa” Foundation under the project code LCF/PR/HR22/52420019. The CNIC is supported by the Instituto de Salud Carlos III (ISCIII), the Ministerio de Ciencia, Innovación Y Universidades (MICIU) and the Pro CNIC Foundation) and is a Severo Ochoa Center of Excellence (grant CEX2020-001041-S funded by MICIU/AEI/10.13039/501100011033).

## Notes

### Competing Interest Statement

The authors have declared no competing interest.

